# Geometric-aware and interpretable deep learning for single-cell batch correction via explicit disentanglement and optimal transport

**DOI:** 10.64898/2026.02.17.706490

**Authors:** Chenghan Jiang, Ren Zheng, Yongle Ji, Shubin Cao, Yanhua Fang, Zhe Wang, Ruoyu Wang, Shanshan Liang, Shuai Tao

**Affiliations:** The Key Laboratory of biomarker high throughput screening and target translation of breast and gastrointestinal tumor, Affiliated Zhongshan Hospital of Dalian University, Dalian, Liaoning, 116001, China; College of Information Engineering, Dalian University, Dalian, Liaoning, 116622, China; National Key Laboratory of Fine Chemical Engineering and Department of Pharmacology in School of Chemical Engineering, Dalian University of Technology, Dalian, Liaoning, 116081, China

**Keywords:** Single-cell RNA sequencing, Batch correction, Explicit disentanglement, Optimal transport, Deep learning

## Abstract

Single-cell RNA sequencing enables high-resolution characterization of cellular heterogeneity, yet integrating datasets from diverse sources remains challenging due to batch effects. Current methods rely on implicit feature disentanglement and and lack geometric constraints, often result in under-correction, over-correction, or compromised biological fidelity. Here, we present iDLC, an interpretable deep learning framework that performs dual-level correction through explicit feature disentanglement and optimal transport–regularized adversarial alignment. iDLC separates biological and technical components within a structured latent space, then leverages high-confidence mutual nearest neighbor pairs to guide geometrically constrained distribution alignment. Systematic evaluation across pancreatic cancer datasets with varying batch effect intensities, multi-source human immune cells, and large-scale cross-species atlases demonstrates that iDLC robustly eliminates complex batch effects while preserving fine-grained cell subtypes, continuous developmental trajectories, and rare populations. The framework scales efficiently to datasets exceeding one million cells and consistently outperforms existing methods in both batch correction and biological conservation metrics. iDLC provides a principled and reliable tool for constructing unified single-cell reference atlases across diverse experimental conditions and biological systems.

## Background

Single-cell RNA sequencing (scRNA-seq) has revolutionized our ability to characterize cellular heterogeneity, offering unprecedented insights into cell types, developmental pathways, and disease mechanisms (Suvà and Tirosh 2019). Large-scale projects like the Human Cell Atlas (Rozenblatt-Rosen et al. 2017) are rapidly generating vast amounts of single-cell data from diverse labs, platforms, and biological samples. To maximize the value of this data, integrating these datasets into a unified, comparable reference atlas has become essential for meaningful biological discovery and clinical translation. However, this integration process is consistently hampered by batch effects—the systematic technical variations introduced by differences in experimental protocols, sequencing platforms, and sample handling (Ye et al. 2023). These variations are often complex and nonlinear, causing cells from different batches to separate artificially in the data, which can severely confound biological interpretation based on cellular similarity (Fröhlich et al. 2019). Currently, three major challenges define the field. First, robustness under strong technical noise: when integrating data with pronounced batch effects—such as multi-platform tumor data—methods must achieve stable correction even when the original biological similarities are heavily distorted, carefully avoiding both under-correction and over-correction (Antonsson and Melsted 2025). Second, preservation of biological fidelity: when processing data containing fine-grained cell subtypes, rare populations, or continuous developmental trajectories, methods must precisely preserve these biologically meaningful structures while removing technical noise, maintaining the natural topology of the cell state space (Pandey and Zafar 2022). Third, specificity in distinguishing sources of variation: in cases where inherent biological differences between datasets (e.g., across species) are far greater than technical batch effects, methods must clearly separate technical noise from genuine biological signals to prevent the erroneous loss of critical information (Lähnemann et al. 2020). Together, these challenges highlight a core need: a new paradigm that can explicitly separate different sources of variation from the start and perform geometry-aware, high-fidelity alignment of distributions.

Unfortunately, current methods have fundamental limitations in addressing these issues. Early linear models like ComBat (Johnson et al. 2007) work well for simple adjustments but struggle with complex nonlinear batch effects. Methods based on mutual nearest neighbors (MNN) and manifold alignment, such as Harmony(Korsunsky et al. 2019) and Scanorama(Hie et al. 2019), identify anchor pairs across batches but can make erroneous matches when batch effects distort the underlying distance metrics. More recently, deep learning approaches like scVI (Lopez et al. 2018), iMAP (Wang et al. 2021) and scDREAMER (Shree et al. 2023) use variational autoencoders or adversarial networks for implicit disentanglement. However, without explicit structural constraints in the latent space, they cannot guarantee a clean separation between biological and technical components, often leading to information leakage and incomplete correction (Yi et al. 2025). Notably, while single-cell foundation models like scGPT (Cui et al. 2024), Geneformer (Theodoris et al. 2023) and scFoundation (Hao et al. 2024) show impressive capabilities in other tasks, they significantly underperform dedicated methods in batch correction (Kömen et al. 2024) —underscoring that model scale alone is not a substitute for targeted architectural design.

To overcome these limitations, we present iDLC (interpretable Dual-Level Correction), a deep learning framework that combines explicit feature disentanglement with optimal transport– regularized adversarial alignment. iDLC tackles the above challenges through a structured two-stage architecture. First, a residual autoencoder with a hard-partitioned latent space explicitly decomposes gene expression into a biological component (encoding cell identity) and a technical component (capturing batch-specific noise). Then, using these purified biological features, iDLC identifies high-confidence MNN pairs to guide an adversarial alignment network, regularized by optimal transport theory. The entropy-regularized Sinkhorn algorithm provides a geometric smoothing constraint, enabling distribution alignment that respects the underlying topology of cell states. Through systematic evaluations on pancreatic cancer data with varying batch effect strengths, multi-source human immune cell datasets, and large-scale cross-species atlases, we demonstrate that iDLC robustly removes complex batch effects while consistently preserving fine-grained subtypes, continuous trajectories, and rare populations. iDLC thus provides a principled, interpretable, and reliable tool for building unified single-cell atlases across diverse experimental and biological contexts.

## Results

### Overview of the iDLC algorithm

The iDLC framework performs batch integration of single-cell data through a structured two-stage deep learning architecture (Figure 1). The core innovation of this algorithm lies in its construction of an end-to-end interpretable deep learning framework, which is achieved by combining explicit feature disentanglement with optimal transport-guided distribution correction.

**Fig. 1.**
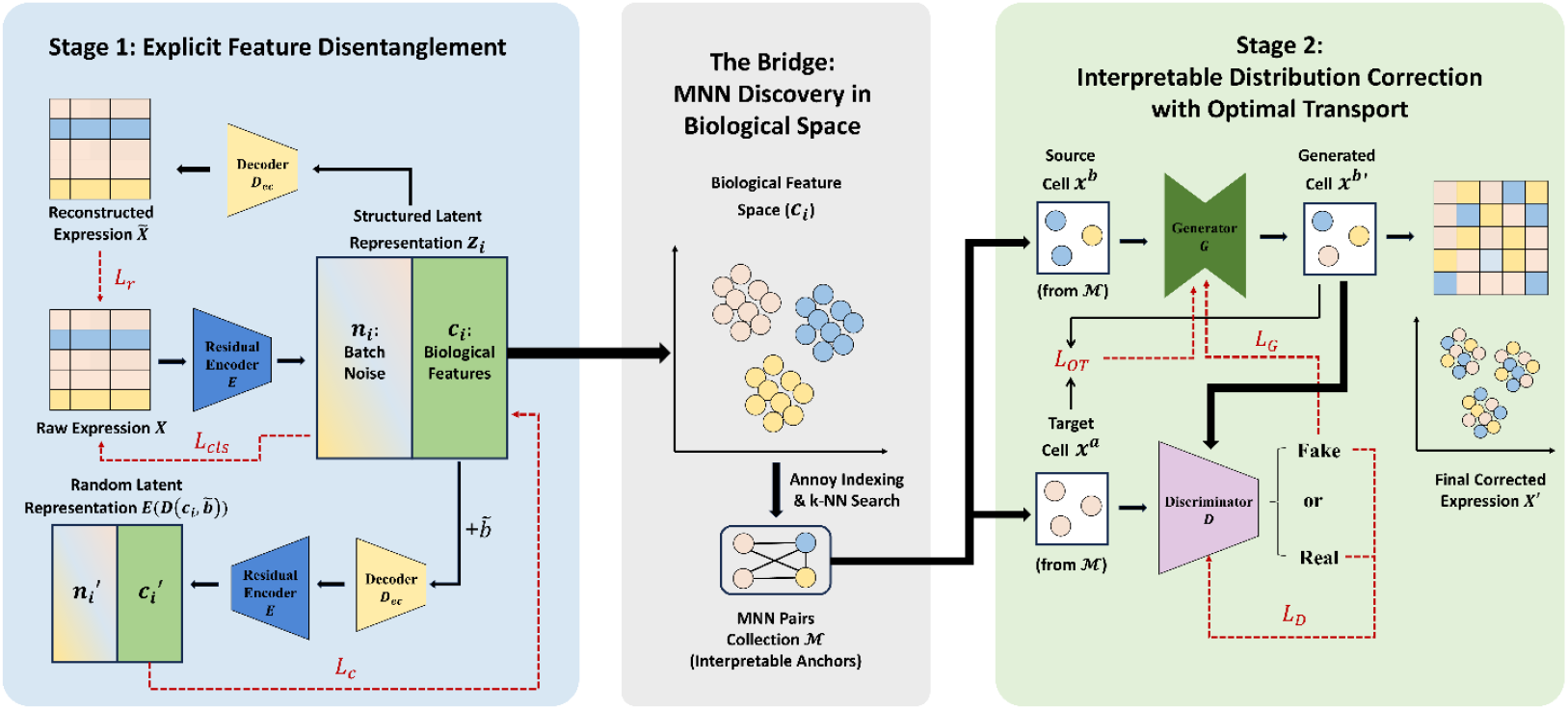
Overview of the iDLC algorithm. Schematic illustration of the two-stage iDLC architecture. The first stage consists of a residual autoencoder that explicitly disentangles gene expression data into biological and batch-noise components in a structured latent space. The second stage employs an optimal transport-regularized generative adversarial network to align distributions across batches while preserving geometric continuity.

In the first stage, the gene expression data *X* is fed into a deep residual autoencoder. The encoder *E*, which consists of multiple fully connected layers and residual blocks, maps the high-dimensional gene expression vector *x*_*i*_ to a structured latent representation *E*(*x*_*i*_). Unlike current deep learning methods that use implicit disentanglement—where the latent representation *E*(*x*_*i*_) and the true batch label *b* are fed into the network with the hope that it learns a purified biological output—iDLC employs explicit disentanglement. This means the latent representation *E*(*x*_*i*_) is explicitly partitioned into two functionally independent subspaces: the first *l* dimensions form the biological feature component *c*_*i*_ and the remaining *k* dimensions form the batch noise component *n*_*i*_ (where *k* is the number of batches). In other words, *E*(*x*_*i*_) = [*c*_*i*_, *n*_*i*_]. This forced split ensures the physical isolation of different variation sources (see Methods).

The decoder *D* reconstructs the gene expression profile *x*_*i*_′ through an inverse transformation. Its training simultaneously optimizes three loss functions: 1) The reconstruction loss *L*_*r*_ = ‖*x*_*i*_′ − *x*_*i*_ ‖ ensures the model accurately captures gene expression patterns; 2) The content consistency loss *L*_*c*_ forces the biological representation *c*_*i*_ to remain batch-invariant by decoding it with a randomly generated batch label 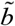; and 3) The batch classification loss *L*_*cls*_ strengthens the noise component *n*_*i*_ ‘s ability to encode batch origin via supervised learning. The synergistic optimization of these three losses enables the model to effectively separate biological signals from technical variation.

In the second stage, the purified biological features from the first stage are used to calculate MNN relationships between batches. Specifically, for any two batches *A* and *B*, high-quality MNN pairs are identified by constructing an Annoy index and performing a k-nearest neighbor search, forming a reliable set ℳ = {(*a, b*) | *a* ∈ *A, b* ∈ *B*}. These high-confidence pairs serve as an interpretable bridge, organically connecting the feature learning of the first stage with the distribution correction of the second.

Next, an optimal transport-regularized generative adversarial network is trained on this MNN pair set ℳ. The generator *G* learns to map the expression profile *x*^*b*^ of a source batch cell to the distribution of the target batch, generating the corrected expression profile 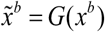. The discriminator *D* distinguishes real cells *x*^*a*^ from corrected cells 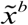 using a Wasserstein distance with gradient penalty strategy. The key innovation here is the introduction of a regularization term based on the Sinkhorn algorithm into the generator’s loss. This optimal transport loss guides the distribution alignment by minimizing the entropy-regularized Wasserstein distance between latent representations, promoting a geometrically smooth, soft assignment. This design allows cells to migrate for batch alignment while preserving local topological structure—which is especially important for protecting dynamic biological processes like continuous developmental trajectories.

The two stages of the framework are tightly connected via the MNN bridge. The explicit disentanglement of the first stage provides a purified biological feature space for obtaining high-quality MNN pairs; these pairs, in turn, supply reliable training samples for the optimal transport-regularized adversarial correction in the second stage. From the original expression data *X*, to the disentangled representation [*c*_*i*_, *n*_*i*_], to the MNN pair set ℳ, and finally to the precise distribution-level correction via the generator *G*, the entire information flow is clear and traceable. This provides strong interpretability guarantees for the final analysis results.

### iDLC robustly integrates pancreatic ductal adenocarcinoma datasets with strong batch effects

To systematically evaluate the robustness of iDLC under different batch effect intensities, we selected two pancreatic ductal adenocarcinoma (PDAC) single-cell transcriptomic datasets that exhibit a clear gradient in technical complexity. The first dataset (mild batch effects) integrates studies by Aadel C et al. (Storrs et al. 2023), Caronni N et al. (Caronni et al. 2023), and Park JK et al. (Park et al. 2024), comprising a total of 202,256 cells. These data were generated under well-controlled experimental conditions with relatively uniform sequencing platforms and protocols, resulting in a low degree of technical variation. The second dataset (strong batch effects) integrates studies by Chen K et al. (Chen et al. 2023), Moncada R et al. (Moncada et al. 2020), and di Magliano MP et al. (Halbrook et al. 2022), containing 128,377 cells. This dataset aggregates samples from different laboratories, employing diverse sequencing technologies and sample preparation protocols, leading to significant and nested batch effects with greater intensity and complexity of technical noise. Both datasets contain the same 12 core pancreatic cell types—including acinar, ductal, and immune cells— providing a consistent benchmark for assessing integration performance within a complex tumor microenvironment (Supplementary Figure S1A, S2A). The central challenge here is to test whether a method can maintain stable correction and biological preservation across this spectrum of technical noise.

On the mild batch-effect dataset, iDLC and all compared methods achieved the basic goal of integration: cells from different batches mixed reasonably well, and major cell type boundaries remained discernible in the visualization (Figure 2A, Supplementary Figure S1B). However, when faced with the strong batch-effect dataset, the limitations of existing methods became apparent (Supplementary Figure S2B). For instance, ComBat, fastMNN, scVI, and scDREAMER showed under-correction, failing to properly integrate T/NK cells across batches. In contrast, iMAP exhibited over-correction, mistakenly aligning epithelial cells with unrelated types like fibroblasts. Harmony and Scanorama, while able to mix batches, disrupted biological continuity—epithelial cells from different batches integrated but appeared in discrete, fragmented clusters. These outcomes highlight the difficulty traditional methods face when technical variation is strong and intertwined with real biological differences.

**Fig. 2.**
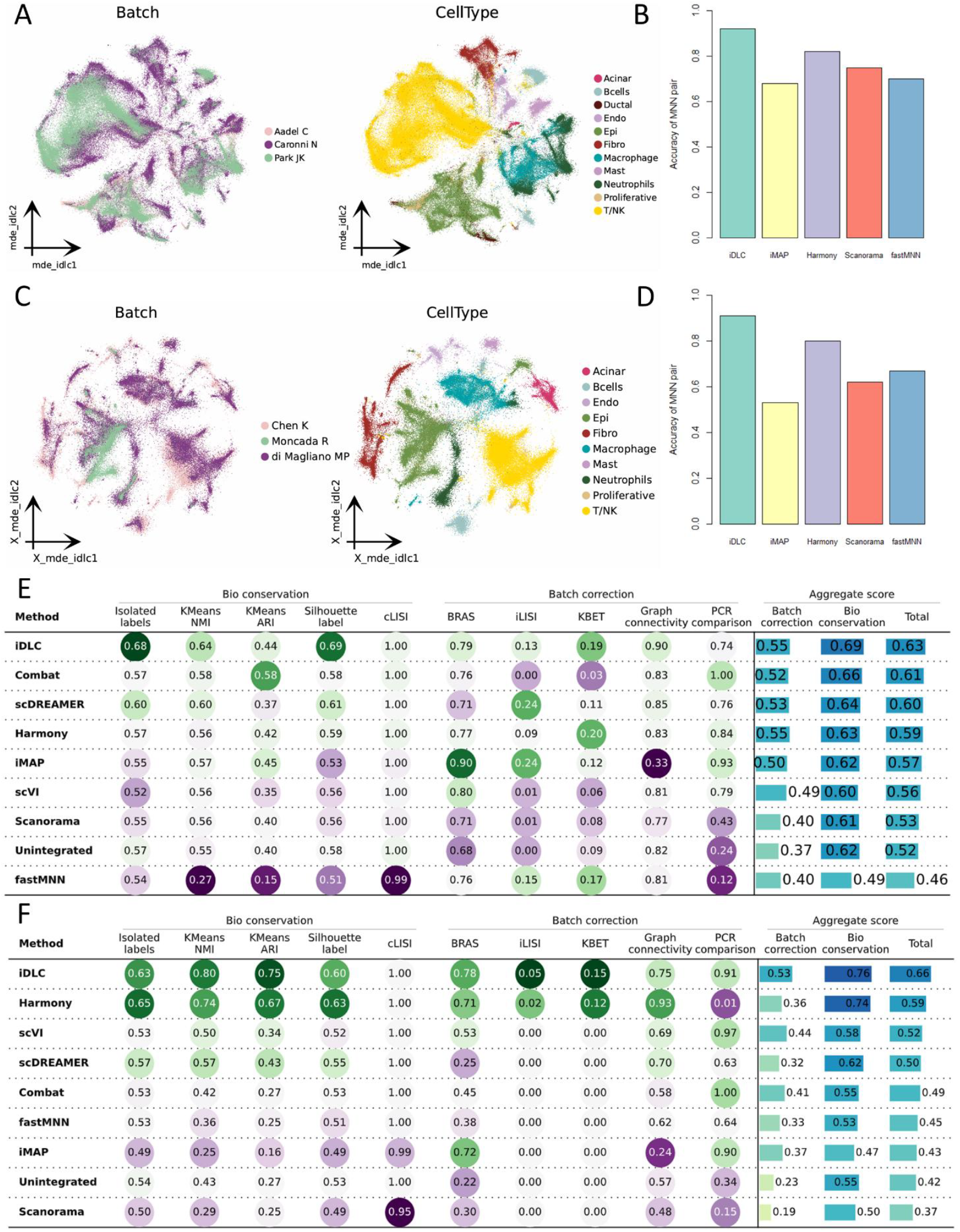
Integration of pancreatic ductal adenocarcinoma datasets with varying batch effect strengths. **(A)** MDE visualization of PDAC data with mild batch effects after integration by iDLC, colored by batch (left) and cell type (right). **(B)** Bar plot comparing the accuracy of mutual nearest neighbor pairs identified by different methods on the mild batch-effect dataset. **(C)** MDE visualization of PDAC data with strong batch effects after iDLC correction, colored by batch (left) and cell type (right). **(D)** MNN pair accuracy across methods under strong batch-effect conditions. **(E, F)** Composite performance scores across ten evaluation metrics for mild (E) and strong (F) batch-effect datasets, summarizing both batch correction and biological conservation.

In contrast, iDLC demonstrated consistent robustness under strong batch effects. Its integrated visualization (Figure 2C) shows thorough mixing of the same cell types across different experimental sources, while all 12 cell types—including hard-to-distinguish proliferating cells and rare populations—remain clearly separated. Both within-cluster compactness and between-type separation were well preserved.

We further investigated the reason behind iDLC’s strong performance. A key hypothesis is that its explicit disentanglement module produces a purified biological feature space, which in turn significantly improves the quality of the MNN anchor pairs used to guide correction. To test this, we quantified the biological accuracy of MNN pairs identified by various methods. On the mild batch-effect data, all methods performed relatively well (Figure 2B). However, on the strong batch-effect data, the quality of MNN pairs from traditional methods dropped substantially when calculated in original or implicitly learned feature spaces (Figure 2D). This means many of the anchor pairs these methods relied on were biologically mismatched, which misleads subsequent correction steps. iDLC, by contrast, still identified high-quality MNN pairs with 89% accuracy even under these challenging conditions, because it builds anchors from an explicitly purified biological feature space. This reliability at the anchor stage explains why iDLC achieves precise yet conservative distribution alignment later on.

Finally, we performed a comprehensive quantitative evaluation using an established metric framework (Luecken et al. 2022). On the mild batch-effect data, all methods reached a usable level of integration, though iDLC consistently achieved the highest overall composite score. Its advantage was especially clear in batch correction metrics, while it also slightly outperformed others in biological conservation (Figure 2E).

On the strong batch-effect dataset, the performance gaps widened considerably. iDLC’s overall score led by a large margin (Figure 2F). It excelled in both correction metrics (highest BRAS, iLISI, kBET) and biological conservation (highest NMI and ARI), showing that it removes technical noise without erasing genuine cell type differences. Other methods revealed specific weaknesses—such as low Graph connectivity or drops in biological scores—quantitatively confirming the under-correction, over-correction, or structural disruption observed visually.

Together, these results demonstrate that iDLC delivers stable and superior integration across varying batch effect strengths, from mild to severe. By using explicit disentanglement to secure reliable biological anchors, and optimal transport to align distributions geometrically, iDLC avoids both under- and over-correction while preserving continuous biological structures. This robustness makes it a generalizable and reliable solution for single-cell data integration in the presence of complex technical variation.

### iDLC accurately integrates human immune cell datasets with fine-grained structures and preserves developmental trajectories

To further assess iDLC’s ability to handle complex biological heterogeneity, we applied it to an integrated human immune cell dataset that combines multiple sources of technical variation with multi-level biological structures. This dataset consists of 33,506 cells from ten independent batches representing different donors. The cells originate from both bone marrow and peripheral blood mononuclear cells (PBMCs) and were sequenced using two distinct protocols—10X Genomics and Smart-seq2 (Supplementary Figure S3A). Among them, 9,581 bone marrow cells are from the study by Oetjen et al. (Oetjen et al. 2018) (3 batches), while the remaining 23,925 PBMCs integrate data from multiple studies including 10X Genomics (Zheng et al. 2017), Freytag et al. (Freytag et al. 2018), Sun et al. (Sun et al. 2019), and Villani et al. (Villani et al. 2017) (totaling 7 batches). This integration task presents several challenges: significant donor-to-donor variation, systematic platform-specific biases, and biological differences due to tissue origin (bone marrow vs. blood). Moreover, the dataset contains two important biological preservation challenges. First, it includes many transcriptionally similar and difficult-to-distinguish cell subtypes, such as CD8^+^ vs. CD4^+^ T cells and CD14^+^ vs. CD16^+^ monocytes, as well as tissue-specific populations found only in bone marrow (e.g., monocyte progenitors and erythroid cells). Second, it encompasses a continuous developmental trajectory from hematopoietic stem and progenitor cells (HSPCs) through erythroid progenitors to mature erythrocytes. Preserving this dynamic continuum across batches is a key test of an integration method’s biological fidelity.

iDLC successfully integrated this highly heterogeneous dataset. In the corrected visualization (Figure 3A), cells from different donors, tissues, and platforms mixed thoroughly according to biological type and formed well-defined clusters. iDLC not only aligned the same cell types across tissues but also cleanly separated key immune subtypes—such as CD8^+^ and CD4^+^ T cells, CD20^+^ and CD10^+^ B cells, and CD14^+^ and CD16^+^ monocytes—into distinct yet neighboring groups. Furthermore, iDLC fully preserved tissue-specific rare populations and clearly maintained the continuous developmental trajectory from HSPCs to erythroid cells, showing that its correction process retains important dynamic biological information.

**Fig. 3.**
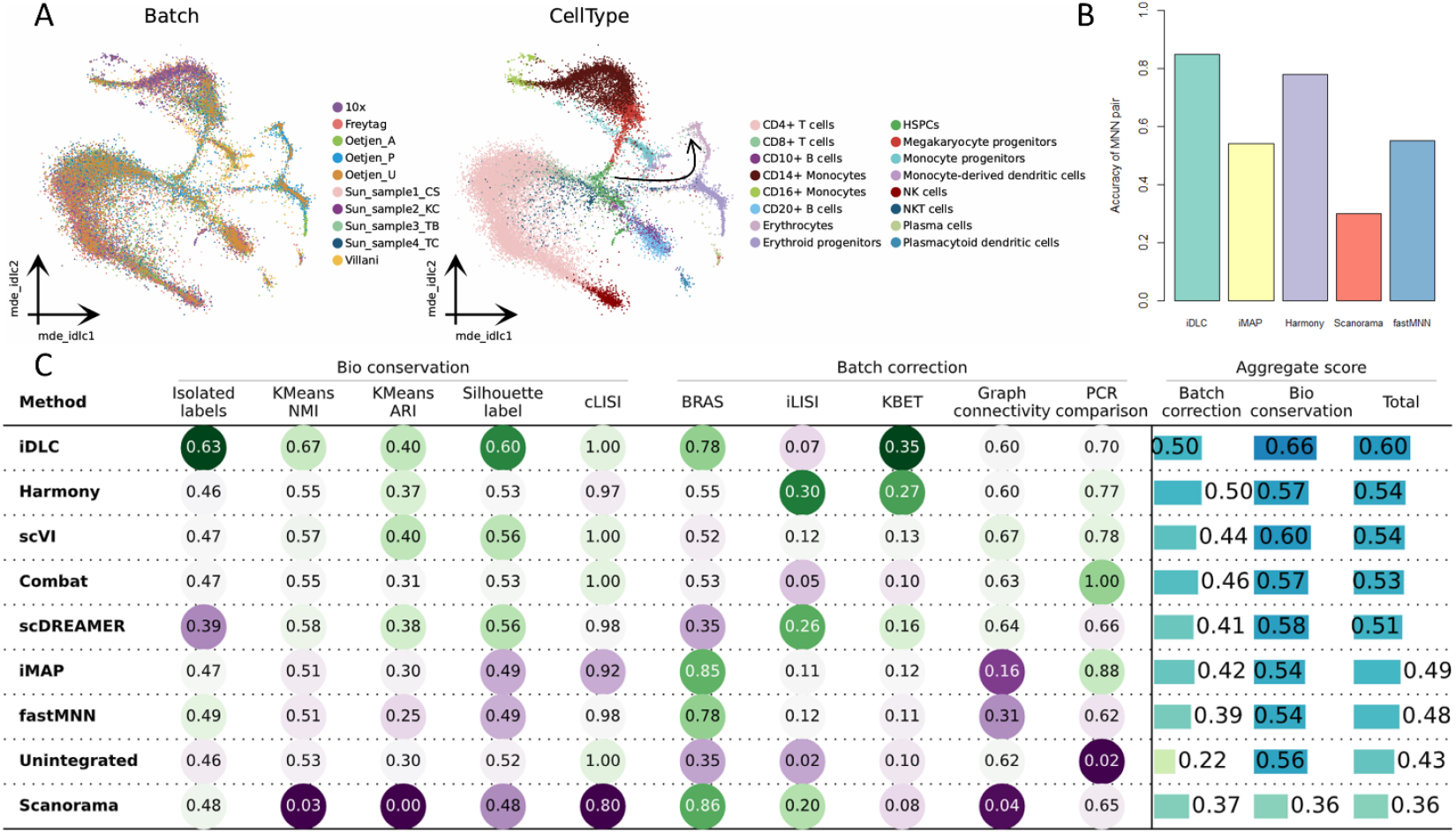
Integration of human immune cell data across donors, tissues, and protocols. **(A)** MDE visualization of integrated human immune data after iDLC correction, colored by batch (left) and annotated cell types/subtypes (right). iDLC preserves fine-grained structures including CD4^+^/CD8^+^ T cells and the erythroid developmental trajectory. **(B)** Accuracy of MNN pairs identified by different MNN-based methods. iDLC achieves the highest accuracy. **(C)** Comprehensive quantitative evaluation using ten metrics and total score. iDLC outperforms other methods, balancing batch removal and biological conservation.

In comparison, existing integration methods revealed various limitations on this complex task (Supplementary Figure S3B). ComBat, Harmony, fastMNN, and scVI displayed under-correction, failing to properly integrate CD4^+^ T cells and CD14^+^ monocytes across batches. Scanorama and scDREAMER exhibited over-correction; Scanorama largely lost the ability to distinguish cell types, while scDREAMER incorrectly merged CD16^+^ monocytes with plasma cells. iMAP, although achieving some mixing, failed to preserve the continuous HSPC–erythroid trajectory—a sign that standard distribution alignment can disrupt biological continuity. Together, these shortcomings highlight common difficulties faced by existing methods when dealing with multiple heterogeneity factors and high-order biological structures simultaneously.

Quantitative analysis supported these observations. On the accuracy of MNN pairs—a metric reflecting the reliability of cross-batch biological correspondences—iDLC significantly outperformed all other MNN-based methods (Figure 3B). This confirms that the explicitly purified biological feature space provides more trustworthy cell-to-cell matches in highly heterogeneous contexts. In the comprehensive performance evaluation, iDLC achieved the highest overall composite score (Figure 3C). Breaking down the scores shows iDLC’s advantage comes from leading across several key sub-metrics.

For biological conservation, iDLC attained the highest Isolated Labels F1, NMI, ARI, and Silhouette label scores, indicating strong preservation of within-cluster compactness, between-cluster separation, and agreement with known biological labels. For batch correction, iDLC earned the highest kBET score, reflecting effective removal of batch differences in local neighborhoods. The low Graph connectivity score of iMAP also aligns with the visual finding that it disrupted developmental continuity.

In summary, this section demonstrates that iDLC can effectively handle the combined influence of multiple heterogeneity factors—donor, platform, tissue, and developmental continuum. Through explicit disentanglement and optimal transport regularization, iDLC removes technical variation while protecting fine-grained subtypes and continuous trajectories, overcoming the typical limitations of existing methods. This makes iDLC a reliable tool for integrating highly heterogeneous single-cell data rich in complex biological structures.

### iDLC enables scalable and robust integration of cross-species single-cell atlases

To test iDLC under an extremely complex and biologically meaningful scenario, we evaluated its performance on a large-scale cross-species integration task. We combined data from the Human Cell Landscape (HCL) (Han et al. 2018) and the Mouse Cell Atlas (MCA) (Han et al. 2020), totaling approximately 933,000 cells. Specifically, HCL contains 599,926 cells spanning 63 human cell types, while MCA contains 333,778 cells covering 52 mouse cell types. Although the two atlases include 97 distinct cell types in total, only 18 are shared between human and mouse. This task presents multiple challenges: the data volume tests algorithmic scalability; the inherent biological differences between species are much larger than typical technical batch effects; and the limited number of shared cell types makes it difficult to identify correct biological correspondences amid vast initial distribution differences (Supplementary Figure S4A).

iDLC successfully performed this atlas-level integration. In the corrected latent space visualization (Figure 4A), cells formed clear clusters based on biological function rather than species origin. Key shared cell types—such as neutrophils, erythroid cells, fetal stromal cells, and oligodendrocytes—showed clear cross-species mixing after integration, with human and mouse cells of the same type accurately aggregating together. This indicates that iDLC effectively separated the confounding factor of species difference and correctly aligned evolutionarily conserved cell states.

**Fig. 4.**
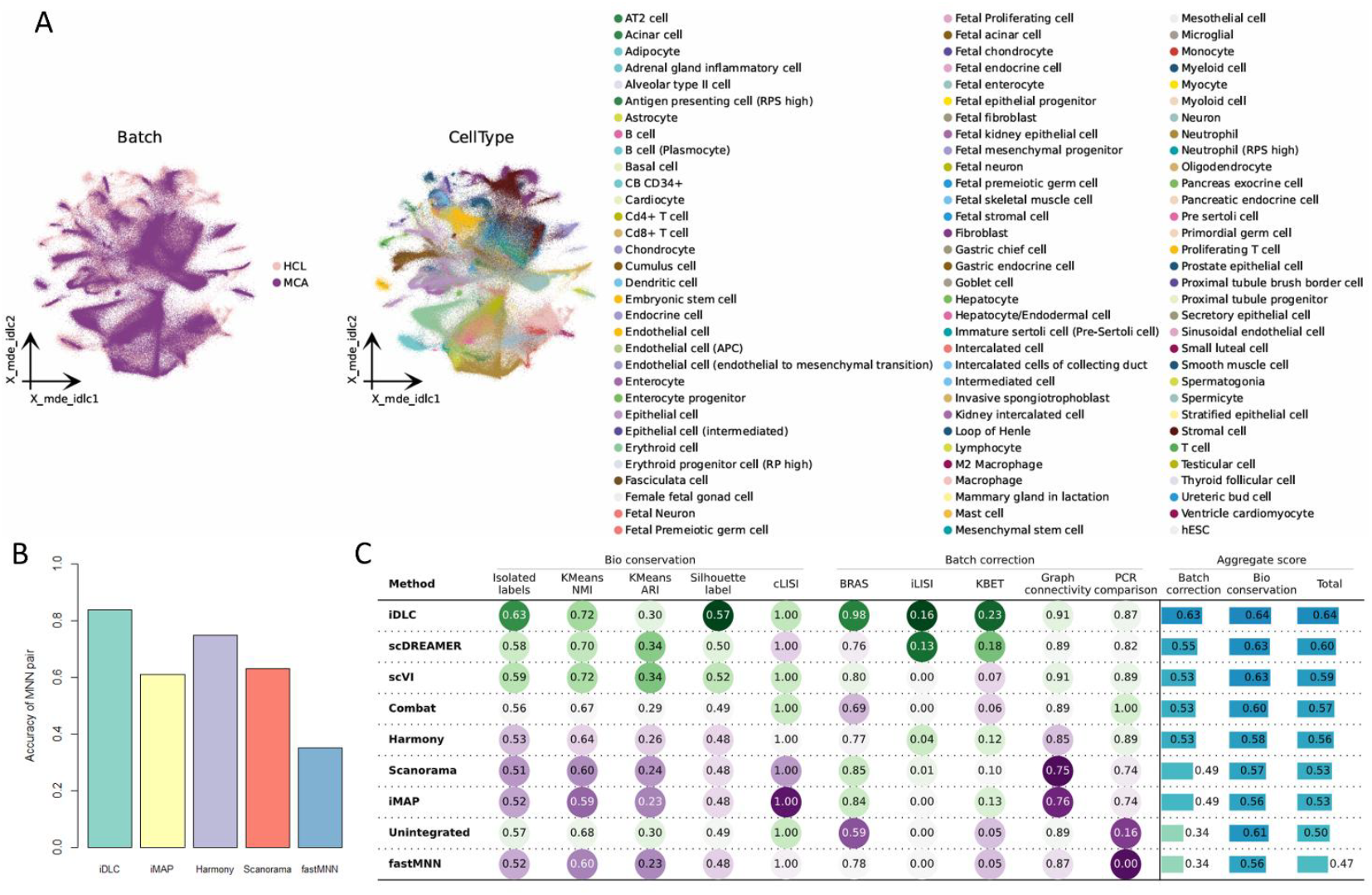
Cross-species integration of human and mouse single-cell atlases. **(A)** MDE visualization of integrated cross-species atlas (∼933k cells) after iDLC correction, colored by species (left) and cell type (right). iDLC aligns shared cell types across species. **(B)** Accuracy of MNN pairs identified by different methods, showing iDLC’s ability to capture conserved biological correspondence. **(C)** Quantitative evaluation of cross-species integration. iDLC achieves the best overall performance among all methods.

On this challenging task, other methods exhibited notable limitations (Supplementary Figure S4B). ComBat and fastMNN showed under-correction, failing to adequately reduce inter-species separation. Scanorama and iMAP displayed over-correction, incorrectly merging different cell types across species. Harmony produced embeddings where similar cell types appeared in disjointed clusters, while scVI and scDREAMER generated fragmented overall distributions. These results suggest that existing methods struggle when biological differences dominate technical variation.

Quantitative analysis further supported these findings. For identifying cross-species biological correspondences, the MNN pair accuracy achieved by iDLC was significantly higher than that of other unsupervised MNN-based methods (Figure 4B). This indicates that even with strong species differences, iDLC’s explicitly disentangled biological features can robustly capture conserved biological similarity.

In the comprehensive evaluation (Figure 4C), iDLC achieved the highest overall composite score among all methods. It performed especially well on batch (species) mixing metrics (BRAS, kBET, iLISI), showing effective integration at both global and local scales. iDLC also ranked first in biological conservation, excelling in metrics such as Isolated Labels F1, NMI, and Silhouette label, confirming its ability to preserve cell type structure and rare populations while removing species-specific bias. Other methods showed various degrees of performance drop or imbalance, quantitatively reflecting the insufficient mixing or structural disruption seen in the visualizations.

In summary, this section demonstrates that iDLC can effectively integrate large-scale cross-species atlas data. The method is scalable to massive datasets and, through explicit disentanglement and optimal transport, can precisely align conserved cell states while separating species-specific features. This overcomes core limitations of existing approaches and provides a powerful, interpretable framework for building unified cross-species atlases and studying evolutionary cell biology.

### Ablation study

To validate the contribution of iDLC’s core design components, we conducted systematic ablation experiments assessing the role of its two key innovations: explicit disentanglement and optimal transport regularization. We compared the full iDLC model against two variants: iDLC without the explicit disentanglement module (iDLC-woED), which replaces the hard-split latent representation with traditional implicit disentanglement; and iDLC without the optimal transport regularization (iDLC-woOT), which removes the optimal transport regularization term from the second-stage generative adversarial network. This allowed us to evaluate how each component influences overall integration performance.

The results across all four benchmark datasets, summarized in Table 1, clearly show that both components are essential for iDLC’s high performance. On the PDAC data with mild batch effects, both variants already showed measurable performance drops. However, in more challenging scenarios—the PDAC data with strong batch effects, the complex immune dataset, and the cross-species integration task—their limitations became much more pronounced.

**Table 1.**
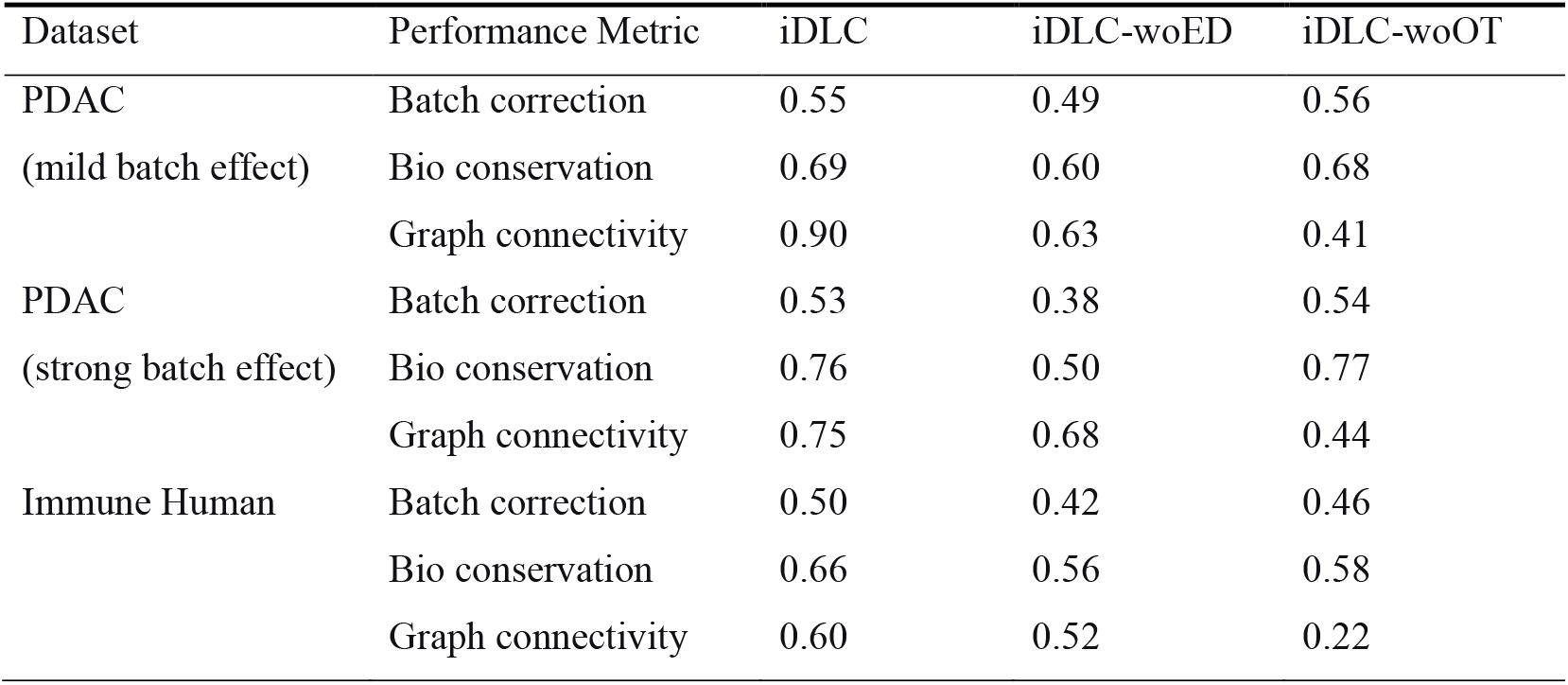
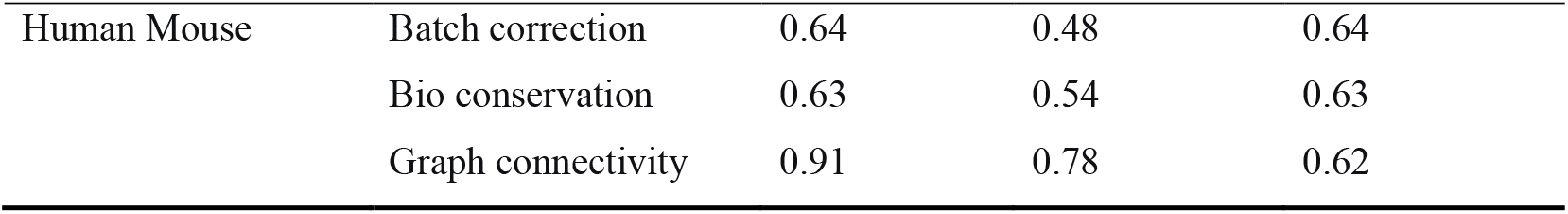
Performance comparison of iDLC and its ablated variants across all benchmark datasets.

Specifically, iDLC-woED (without explicit disentanglement) exhibited noticeable under-correction and biological structure confusion on datasets with strong batch effects and cross-species differences. Its biological conservation score, for example, fell sharply from 0.76 to 0.50 on the strong batch-effect PDAC data. This confirms that explicit disentanglement is critical for cleanly separating biological signals from technical noise at the source, which in turn enables reliable anchor identification.

Meanwhile, iDLC-woOT (without optimal transport regularization) maintained batch correction scores close to the full model, but its Graph connectivity score decreased substantially across all datasets—most notably from 0.60 to 0.22 on the immune data. This decline reflects a reduced ability to preserve the continuity of cell states and local topological relationships, underscoring the importance of optimal transport in maintaining dynamic biological processes such as developmental trajectories.

Taken together, these ablation experiments demonstrate that explicit disentanglement and optimal transport regularization work synergistically within the iDLC framework. Explicit disentanglement isolates biological and technical factors at the feature level, providing a reliable foundation for cross-batch alignment. Optimal transport regularization then ensures that the distribution alignment process respects geometric continuity, preventing the disruption of inherent biological structures. The absence of either component leads to measurable performance degradation, particularly in difficult integration scenarios.

## Discussion

In this study, we introduced iDLC, an interpretable deep learning framework for single-cell data integration that combines explicit feature disentanglement with optimal transport–regularized adversarial alignment. Our design directly addresses several key challenges in current batch correction methods. First, unlike mainstream approaches that rely on loss functions to implicitly separate signals in a “black-box” manner(Yi et al. 2025), iDLC introduces a structured explicit disentanglement mechanism. By forcing a hard split of the latent space in a residual autoencoder, we ensure the physical isolation of biological and technical variation at the architectural level. This not only makes the separation more reliable and complete, but also traceable—significantly improving interpretability. Thanks to this explicit disentanglement, iDLC extracts a purified biological feature space, which consistently yields MNN pairs with higher biological accuracy across all tested scenarios, from mild to strong batch effects and even in cross-species integration. These high-quality correspondences then provide a trustworthy anchor set for the subsequent distribution alignment. Second, to address the common issue where distribution alignment disrupts the topology of cell state space, iDLC innovatively incorporates optimal transport theory as a regularization constraint within the adversarial learning stage. The soft assignment and geometrically smooth matching achieved via the Sinkhorn algorithm overcome the limitations of traditional hard alignment strategies. This provides crucial mathematical assurance for preserving dynamic biological structures such as continuous developmental trajectories. Across a series of realistic and challenging tasks—pancreatic cancer data with gradient batch effects, multi-source human immune data, and large-scale cross-species atlas integration—iDLC consistently demonstrated superior robustness and precision. Ablation studies further confirmed that both the explicit disentanglement module and the optimal transport regularization are indispensable for iDLC’s performance. Each component contributes critically to maintaining biological fidelity and correction stability, especially in difficult integration scenarios. Overall, the evidence indicates that iDLC achieves one of the best current balances between removing batch effects and preserving fine-grained biological structures.

Despite its strong performance, iDLC has some limitations that point toward valuable future research directions. At the model level, the division of latent space dimensions in the explicit disentanglement module still relies on prior knowledge or heuristic settings. Future work could explore adaptive dimension inference mechanisms to improve generalizability. Additionally, while entropy regularization accelerates optimal transport computation, its computational and memory overhead can still be challenging for extremely large-scale data (e.g., tens of millions of cells). Developing more efficient approximation algorithms or hierarchical processing strategies will be important for engineering optimization. In terms of methodological extensions, a natural next step is applying this explicit disentanglement paradigm to multimodal single-cell data integration (e.g., scRNA-seq with ATAC-seq or proteomics) (Ranzoni et al. 2021). The goal would be to separate cross-omics biological modules from modality-specific technical noise (Suo et al. 2022). Moreover, although optimal transport regularization significantly improves the preservation of continuous structures, future architectures might incorporate dynamical priors—such as neural ordinary differential equations—to more finely model cell state transitions (Li et al. 2024). Finally, developing adaptive hyperparameter optimization strategies and automated batch structure inference modules would enhance iDLC’s robustness and ease of use when dealing with data of unknown or nested batch complexity (Argelaguet et al. 2021; Jiang et al. 2024).

The principles established by iDLC also have broader implications for biomedical research. For fundamental science, iDLC provides a powerful tool for building high-quality, comparable single-cell reference atlases across conditions, platforms, and even species (Baião et al. 2025). This directly supports the refinement of cell atlases, deeper analysis of disease heterogeneity, and studies of evolutionary conservation. Its clear disentanglement capability also makes it possible to isolate and quantify the impact of specific technical factors—such as different sample preparation protocols—offering an objective computational means to standardize and optimize experimental workflows. In translational and clinical contexts, iDLC can efficiently integrate patient data collected across multiple centers or time points. This enhances the detection of rare cell subtypes, improves the reliability of biomarker discovery, and supports more robust inference of cell state trajectories. More broadly, by building interpretability directly into the architecture—rather than relying on post-hoc explanations—iDLC helps advance single-cell data integration from implicit fitting toward explicit, principled analysis. This paradigm shift may inform the development of more reliable and transparent computational tools across biomedical data science.

## Methods

### The iDLC model

The core idea of the iDLC framework draws inspiration from the concept of explicit feature disentanglement in the field of computer vision. Just as image representations can be explicitly separated into independent factors such as content (e.g., object shape) and style (e.g., texture, lighting), iDLC introduces this paradigm for the first time to single-cell transcriptomic data integration: we structurally disentangle the gene expression representation into a biological component encoding the intrinsic cell identity and a batch-noise component capturing technical variation. This design fundamentally breaks through the implicit disentanglement paradigm relied upon by current mainstream methods—the latter only performs ambiguous signal separation in an uninterpretable latent space. Through this explicit disentanglement architecture, iDLC ensures the physical isolation of biological signals from technical noise at the model level, thereby significantly enhancing the interpretability of the correction process and providing a more robust solution for handling complex batch effects.

Our model is built upon recent advances in deep learning, particularly ideas from deep generative models and adversarial learning. Unlike traditional implicit learning approaches, iDLC introduces an explicit disentanglement mechanism that enforces feature separation at the architectural level, thereby ensuring the interpretability of the entire correction process. The framework consists of two sequentially connected stages: an explicit feature disentanglement stage based on a residual autoencoder, and an adversarial distribution alignment stage guided by high-quality cell pairs.

During model construction, we assume that the gene expression profile of a single cell, *x* ∈ ℝ^*g*^ (where *g* represents the number of highly variable genes), is jointly determined by two independent latent factors: biological content *c* ∈ ℝ^*l*^ and batch-specific technical noise *n* ∈ ℝ^*k*^. Here, *l* denotes the dimensionality of the biological representation, and *k* represents the number of batches. This factorized assumption allows us to model and process biological signals and technical noise separately.

Unlike probabilistic generative models such as Variational Autoencoders (VAEs), iDLC employs a deterministic encoder-decoder architecture, enforcing a hard split at the latent representation level to ensure physical isolation of variations from different sources in the feature space. This approach not only provides better model interpretability but also enables more precise control over the feature disentanglement process.

### Explicit Disentanglement with Residual Autoencoder

The first stage of the iDLC framework employs an explicit disentanglement architecture based on a residual autoencoder, aiming to achieve the structured separation of biological variation and batch effects in single-cell transcriptomic data. The core innovation of this method lies in the physically isolated latent representation space through a hard split, separating technical noise from biological content at the feature level, thereby ensuring the reliability and interpretability of subsequent analyses.

We assume that the gene expression profile of each cell is jointly determined by two independent latent factors: biological content reflecting cell type, state, and function, and batch noise originating from technical factors such as experimental conditions and sequencing platforms. This assumption is based on the generative mechanism of single-cell sequencing data, where observed gene expression differences can be decomposed into true biological differences and systematic errors introduced by technology.

The deep residual autoencoder deployed in this stage consists of an encoder *E* and a decoder *D*. The encoder maps the input gene expression vector *x* ∈ ℝ ^*g*^ to a structured latent representation *E*(*x*) ∈ ℝ ^(*l+k*)^, where *g* is the number of highly variable genes, *l* is the dimensionality of biological features, and *k* is the number of batches. This representation is explicitly partitioned into two functionally independent subspaces: the first *l* dimensions constitute the biological feature component *c*, specifically capturing cell identity and state information; the remaining *k* dimensions constitute the batch noise component *n*, specifically encoding technical variation information.

During training, we introduce an innovative content consistency verification mechanism. Specifically, for an input cell *x*, its explicitly disentangled feature representation [*c, n*] = *E*(*x*) is first obtained via the encoder. Subsequently, random batch noise 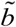 is generated, where 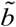 is a randomly generated one-hot vector representing a random batch label. This random noise is combined with the original biological feature *c* and then passed through the decoder to reconstruct a pseudo-cell 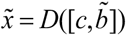. Finally, the pseudo-cell is input into the encoder again to extract its corresponding biological feature 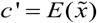.

To ensure effective feature disentanglement, we designed three complementary loss functions. The reconstruction loss ensures the model accurately captures gene expression patterns:

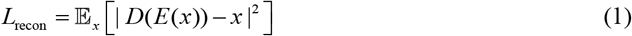

where *x* denotes the input gene expression vector and *D*(*E*(*x*)) denotes the reconstructed expression vector. This loss guarantees the basic functionality of the autoencoder by minimizing reconstruction error.

The content consistency loss forces biological features to remain stable across different batch conditions:

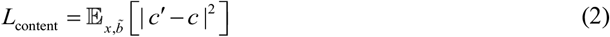

where *c* denotes the biological feature of the original cell and *c* ′ denotes the biological feature of the pseudo-cell. This loss ensures the batch invariance of the biological representation by constraining feature consistency.

The batch classification loss ensures, through supervised learning, that the noise component can effectively distinguish batch origins:

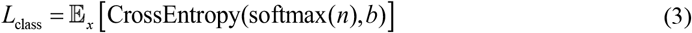

where *n* denotes the batch noise component output by the encoder and *b* denotes the true batch label. This loss function forces the batch noise component *n* to contain sufficient information to accurately predict the cell’s batch origin by minimizing the cross-entropy error of batch prediction, thereby ensuring it specifically captures technical variation rather than biological signals.

The overall optimization objective is a weighted combination of the three loss functions:

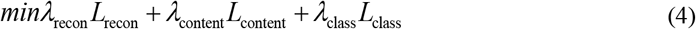

where *λ*_recon_, *λ*_content_ and *λ*_class_ are hyperparameters used to balance the relative importance of the different optimization objectives.

Through this multi-objective optimization strategy, the encoder learns a pure, batch-invariant biological feature representation. These features provide a reliable foundation for the distribution alignment in the second stage while ensuring the transparency and interpretability of the entire correction process.

### MNN Pair Identification as the Integration Bridge

After obtaining the purified biological feature representation generated in the first stage, we need to construct a reliable bridge connecting the disentangled features with the subsequent distribution alignment. This bridge consists of MNN pairs, with the core objective of discovering high-confidence cross-batch cell correspondences in the disentangled biological feature space, thereby providing high-quality, biologically credible training sample pairs for the adversarial correction in the second stage.

Specifically, for batch *A* and batch *B*, we first normalize their biological features:

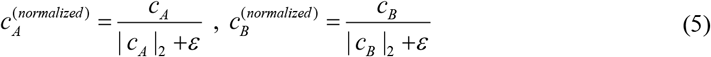

where *c*_*A*_ and *c*_*B*_ represent the biological features of cells in batch *A* and batch *B* respectively, and *ε* is a constant set for numerical stability.

Next, we use an efficient approximate nearest neighbor algorithm (e.g., Annoy) to construct MNN pairs. For each cell *a* in batch *A*, we search for its *K* nearest neighbor cells in batch *B*; similarly, for each cell *b* in batch *B* we search for its *K* nearest neighbor cells in batch *A*. A cell pair (*a, b*) is identified as a valid MNN pair if and only if cell *a* is among the *K* nearest neighbors of cell *b*, and cell *b* is also among the *K* nearest neighbors of cell *a* :

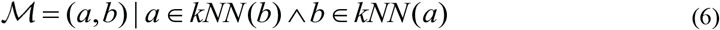

where *kNN* (·) denotes the set of *K* nearest neighbor cells in the target batch.

Compared to traditional MNN methods, a key advantage of iDLC lies in its utilization of the purified biological feature space after explicit disentanglement. Since this space has largely removed the interference of technical noise through the hard split in the first stage, the inter-cellular distances calculated in this space more accurately reflect biological similarity. Consequently, the identified MNN pairs exhibit higher biological relevance and accuracy, providing reliable supervisory signals for the next stage. This collection of high-quality cell correspondences serves as interpretable anchors, guiding the optimal transport generative adversarial network in the second stage to learn how to align the distributions of different batches.

### Optimal Transport-guided Adversarial Correction

The second stage of the iDLC framework achieves precise and geometry-aware distribution alignment through an optimal transport-regularized generative adversarial network. This stage uses the MNN pair set ℳ as supervisory signals to train a generator network that maps the expression profiles of source batch cells into the distribution of the reference batch. The core innovation lies in integrating optimal transport theory as a key regularization term into the adversarial training, ensuring that the distribution alignment process not only eliminates batch differences but also preserves the continuity and local topology of the cell state space.

The constructed generative adversarial network consists of a generator *G* and a discriminator *D*. The generator’s core function is to map the expression profiles of source batch cells to the distribution space of the reference batch. Its design employs a neural network architecture with residual connections:

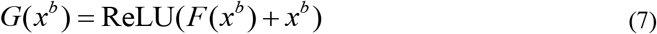

where *x*^*b*^ denotes the expression profile of cell *b* from batch *B* in an MNN pair, and *F* is a deep autoencoder network. This residual design ensures the stability of the generation process.

The discriminator is responsible for distinguishing real reference batch cells from corrected cells produced by the generator, constructed using a multilayer perceptron:

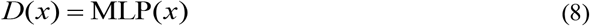

It captures batch-specific distribution features through stacked fully connected layers.

The optimization objective for adversarial training is based on the Wasserstein GAN with Gradient Penalty (WGAN-GP) framework, ensuring training stability and convergence:

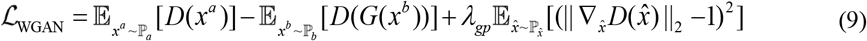

where ℙ _*a*_ and ℙ _*b*_ denote the data distributions of reference batch *A* and source batch *B* respectively, and *λ*_*gp*_ is the gradient penalty coefficient.

The key innovation lies in the enhancement of generator training. We introduce a regularization term based on optimal transport distance into its loss function, encouraging the generator not only to fool the discriminator but also to align the distribution of generated data with the target distribution in a geometrically optimal manner:

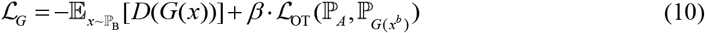

where *β* is a hyperparameter controlling the strength of optimal transport regularization.

The optimal transport regularization term ℒ_OT_ is computed using the entropy-regularized Sinkhorn algorithm. For a batch of real reference samples 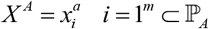 and corresponding generated samples 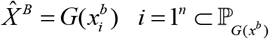 from MNN pairs, we compute their cost matrix *C* ∈ ℝ^*m*×*n*^. In iDLC, to ensure optimal transport alignment is based on biological content, we use the biological latent representations extracted by the first-stage autoencoder to calculate the cost:

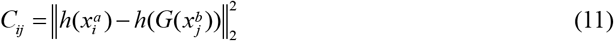

where *h*(·) = Encoder_bio_ (·) is the part of the autoencoder that extracts the biological latent representation.

The entropy-regularized optimal transport problem is defined as:

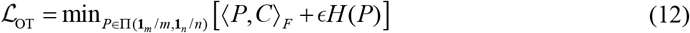

where Π(·) is the set of coupling matrices satisfying uniform marginal distributions, 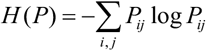 is the entropy term, and ε is the regularization coefficient. The introduction of entropy regularization makes the solution efficient and stable, producing a soft-assignment, probabilistic transport plan *P*, which better reflects the geometric relationship between distributions and is more robust to noise compared to hard matching.

This problem is solved quickly via the Sinkhorn iterative algorithm. Ultimately, the optimal transport loss ℒ_OT_ is this minimal transport cost. By incorporating this loss as a regularization term, the generator *G* is guided to generate data that are not only judged as real by the discriminator but also minimize the transport cost (i.e., the degree of geometric deformation in the biological feature space) from source batch *B* to reference batch *A*.

During training, we sample batch data 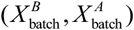 from the MNN pair set and alternately optimize the discriminator *D* and generator *G*. After training is completed, the generator *G* is applied to the entire cell expression matrix *X* ^*B*^ of source batch *B* to obtain the batch-corrected data:

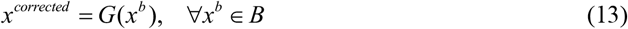

In summary, the second stage of iDLC is an optimal transport-regularized adversarial distribution alignment process. Leveraging the pure biological features provided by the first stage and the high-quality MNN pairs based thereon, it achieves geometrically smooth alignment of the distribution of source batch *B* to reference batch *A* through optimal transport-regularized GAN training, while preserving key biological structures. The entire framework, from explicit disentanglement to optimal transport alignment, forms a logically rigorous and highly interpretable complete pipeline.

### Implementation details

In terms of model implementation, the iDLC framework is built on PyTorch and contains two core components: the residual autoencoder for explicit feature disentanglement and the generative adversarial network for distribution alignment. The residual autoencoder employs a deep encoder-decoder architecture (Figure 5A). The encoder pathway maps from the input dimension *g* to 1024 dimensions via a fully connected layer, followed by a batch normalization layer and ReLU activation function. It then sequentially passes through two residual blocks(He et al. 2016) with dimensions of 1024 and 512, ultimately outputting a structured latent representation with a dimensionality equal to the sum of *l* and *k*. The decoder pathway starts with this latent representation and reconstructs the gene expression profile through a symmetric structure, sequentially passing through residual blocks of 512 and 1024 dimensions, with the final output dimension restored to *g*. The residual blocks within the network (Figure 5B) adopt a bottleneck design with identity mapping, containing two fully connected layers, each followed by batch normalization. The output of the main pathway is added to the skip connection before passing through a ReLU activation, ensuring stable gradient propagation in deep networks.

**Fig. 5.**
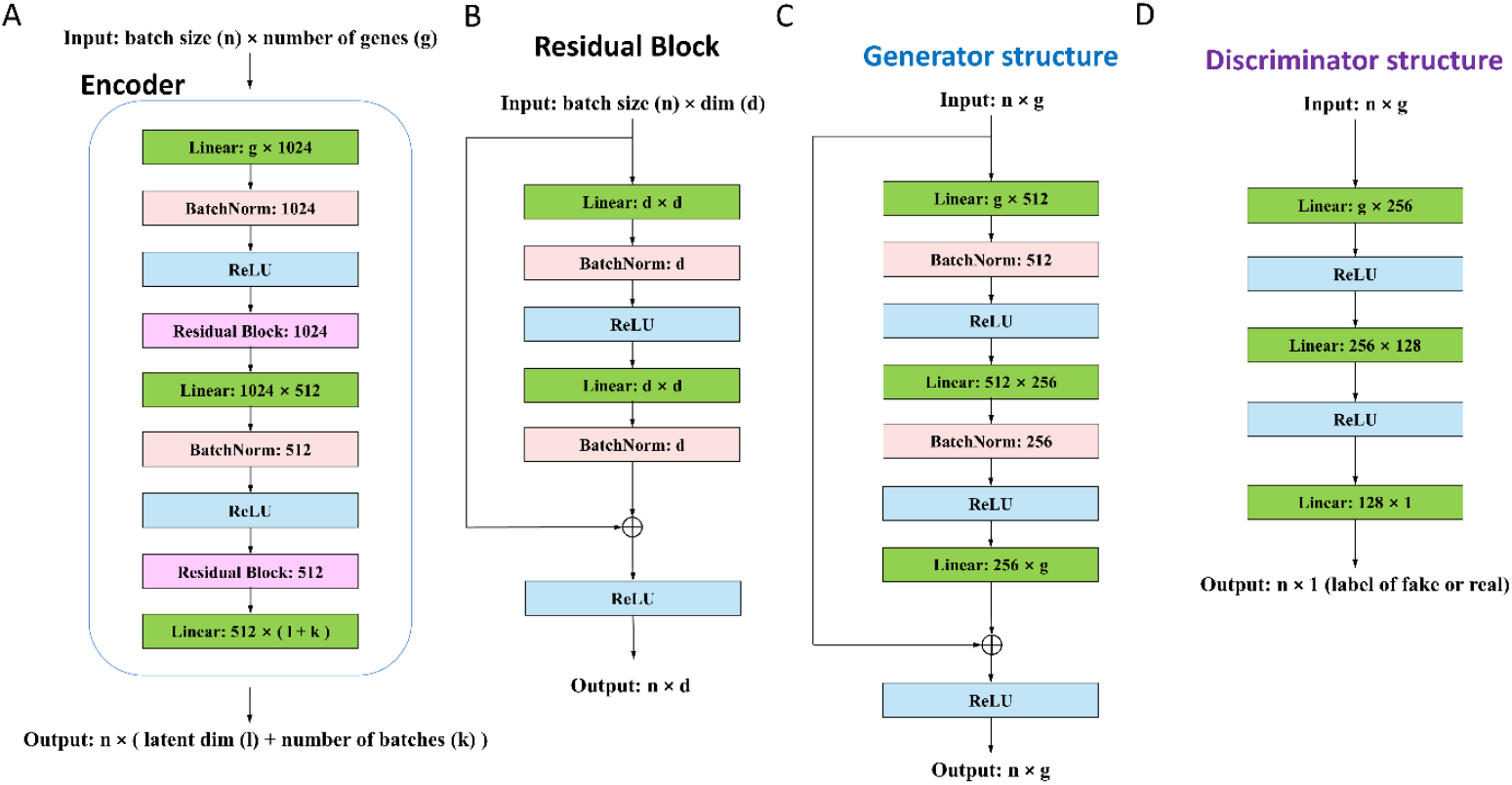
Neural network architectures of the iDLC framework. **(A)** The residual autoencoder for explicit feature disentanglement, showing encoder and decoder pathways with dimensionality transitions. **(B)** Detailed structure of a residual block with skip connections for stable gradient flow. **(C)** The GAN generator with residual connections for distribution transformation. **(D)** The feedforward discriminator network for batch distribution discrimination.

In the generative adversarial network, the generator (Figure 5C) employs an architecture with residual connections. The input data sequentially passes through fully connected layers of 512 and 256 dimensions, each followed by batch normalization and ReLU activation. The final output layer ensures the learning of a distribution transformation rather than complete reconstruction via a ReLU residual connection. The discriminator (Figure 5D) is a concise feedforward network. The input data sequentially passes through fully connected layers of 256 and 128 dimensions, both using ReLU activation, ultimately outputting a single-dimensional discrimination result.

Autoencoder training employs a multi-objective loss function, where the reconstruction loss weight is 1.0, and the content consistency loss and batch classification loss weights are both 0.5. The Adam optimizer with a learning rate of 0.001 is used, along with weight decay of 1e-5 and a learning rate decay strategy.

The adversarial training phase adopts the optimal transport-WGAN-GP framework. The gradient penalty coefficient is 10. The learning rates for both the discriminator and generator are 0.0001, with one generator update corresponding to every five discriminator updates.

All experiments were completed on a Linux server equipped with an NVIDIA Tesla V100S-PCIE-32GB GPU. The batch size was uniformly set to 256. MNN pair identification utilized the Annoy library for efficient approximate nearest neighbor search (K=20). The framework automatically selects the batch with the largest variance as the reference anchor point, enabling flexible handling of different numbers of batches.

### Preprocessing scRNA-seq datasets

Preprocessing of single-cell RNA sequencing data followed the standard Scanpy analysis pipeline. Strict quality control was first performed to filter out low-quality cells and genes, retaining cells expressing at least 200 genes and genes expressed in at least 3 cells. Subsequently, library size normalization was performed for each cell’s gene expression, targeting a total count of 10,000, followed by a log(1+x) transformation on the normalized data to stabilize variance. Selection of highly variable genes employed the Seurat v3 method, considering the influence of batch effects, jointly screening 2000 highly variable genes from all batches for subsequent analysis. Only genes detected in all batches were considered, ensuring the reliability of cross-batch comparisons. The preprocessed gene expression matrix served as the input for the iDLC framework, where each row represents a cell and each column represents the expression value of a highly variable gene.

### Metrics for the evaluation of integration performance

To systematically evaluate the performance of iDLC and competing integration methods, we employed a comprehensive set of ten established metrics that assess two critical aspects of data integration: biological conservation and batch effect correction. All metrics were computed based on the low-dimensional embeddings generated by each integration method to ensure fair comparison. The metrics are categorized as follows:

Biological conservation metrics: This category of metrics evaluates how well an integration method preserves genuine biological variation after removing technical batch effects.

- **Isolated Labels F1 Score** measures the ability to preserve rare cell types. For each rare cell type present in the fewest batches, we calculate the F1 score by treating the rare type as one cluster and all other cells as another cluster:

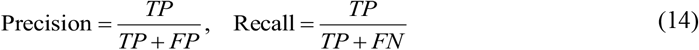

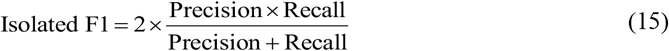

where TP, FP, and FN represent true positives, false positives, and false negatives, respectively.
- **Normalized Mutual Information (NMI)** quantifies the agreement between clustering results and true cell type labels:

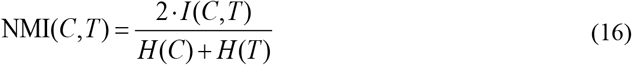

where *I* (*C,T*) represents the mutual information between clustering assignment *C* and true labels *T*, while *H* (·) denotes entropy.
- **Adjusted Rand Index (ARI)** assesses the similarity between two data clusterings by considering all pairs of samples:

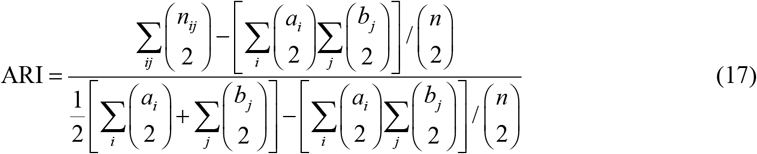

where *n*_*ij*_ denotes the number of cells in cluster *i* of the first clustering and cluster *j* of the second clustering, with *a*_*i*_ = Σ _*j*_ *n*_*ij*_, *b*_*i*_ = Σ_*i*_ *n*_*ij*_, and *n* representing the total number of cells.
- **Cell-type Silhouette Width** measures the compactness of cell types and their separation from other types. For each cell *i* :

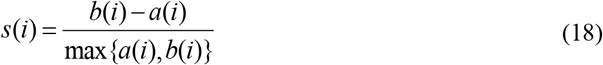

where *a*(*i*) is the average distance from cell *i* to other cells in the same cluster, and *b*(*i*) is the average distance from cell *i* to cells in the nearest neighboring cluster. The final metric is computed as the average over all cells.
- **Cell-type Local Inverse Simpson’s Index (cLISI)** evaluates the purity of cell-type neighborhoods in the integrated space:

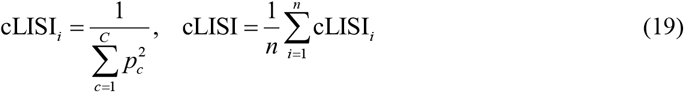

where *p*_*c*_ represents the proportion of cells of type *c* in the local neighborhood of cell *i*. Batch correction metrics: This category quantifies how effectively technical batch effects have been removed while preserving biological signals.
- **Batch-related Axis Score (BRAS)** measures the presence of batch-related axes in the integrated data:

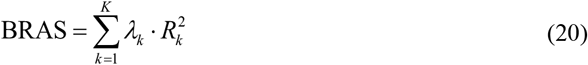

where *λ*_*k*_ is the variance explained by the *k*-th principal component and 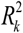 is the coefficient of determination from regressing the principal component on batch labels.
- **Batch Local Inverse Simpson’s Index (iLISI)** assesses the mixing of batches in local neighborhoods:

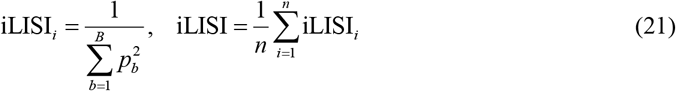

where *p*_*b*_ is the proportion of cells from batch *b* in the local neighborhood of cell *i*.
- **k-nearest neighbor Batch Effect Test (kBET)** evaluates whether the local batch composition around each cell matches the global batch distribution:

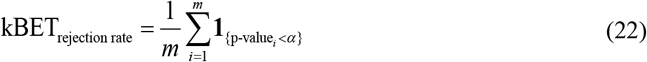

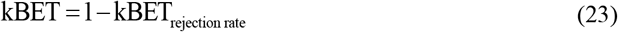

where the test is performed on *m* randomly sampled cells, with p-values computed using chi-square tests for local batch distributions.
- **Graph Connectivity** measures whether cells from the same cell type form connected components in the k-nearest neighbor graph:

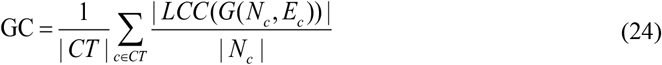

where *CT* represents the set of cell types, *LCC*(·) denotes the largest connected component, and *G*(*N*_*c*_, *E*_*c*_) is the kNN subgraph for cell type *c*.
- **Principal Component Regression (PCR)** quantifies the amount of variance in principal components explained by batch information:

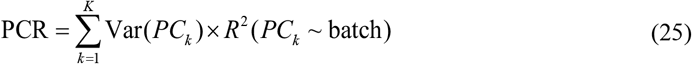

where Var(*PC*_*k*_) is the variance explained by the *k*-th principal component.

Composite scoring: To provide a comprehensive assessment of integration performance, we computed composite scores that aggregate performance across multiple metrics without normalization. The scoring procedure was conducted as follows:

Biological Conservation Score was calculated as the arithmetic mean of all five biological conservation metrics:

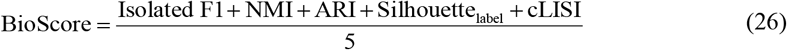

Batch Correction Score was calculated as the arithmetic mean of all five batch correction metrics:

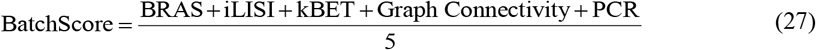

Total Score was then computed as the average of the Biological Conservation Score and Batch Correction Score:

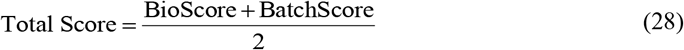

This scoring framework provides a balanced evaluation of each method’s ability to simultaneously preserve biological signals and remove technical artifacts, with higher scores indicating better overall integration performance across both critical dimensions.

### Competing integration methods

We compared our method with seven state-of-the-art integration methods: Combat(Johnson et al. 2007), Harmony(Korsunsky et al. 2019), fastMNN(Haghverdi et al. 2018), Scanaroma(Hie et al. 2019), scVI(Lopez et al. 2018), iMAP(Wang et al. 2021) and scDREAMER(Shree et al. 2023). Detailed information regarding the running configurations of these methods is provided in Supplementary Table S1.

## Supplementary Information

For details, refer to the supplementary Information.

**Fig. S1.**
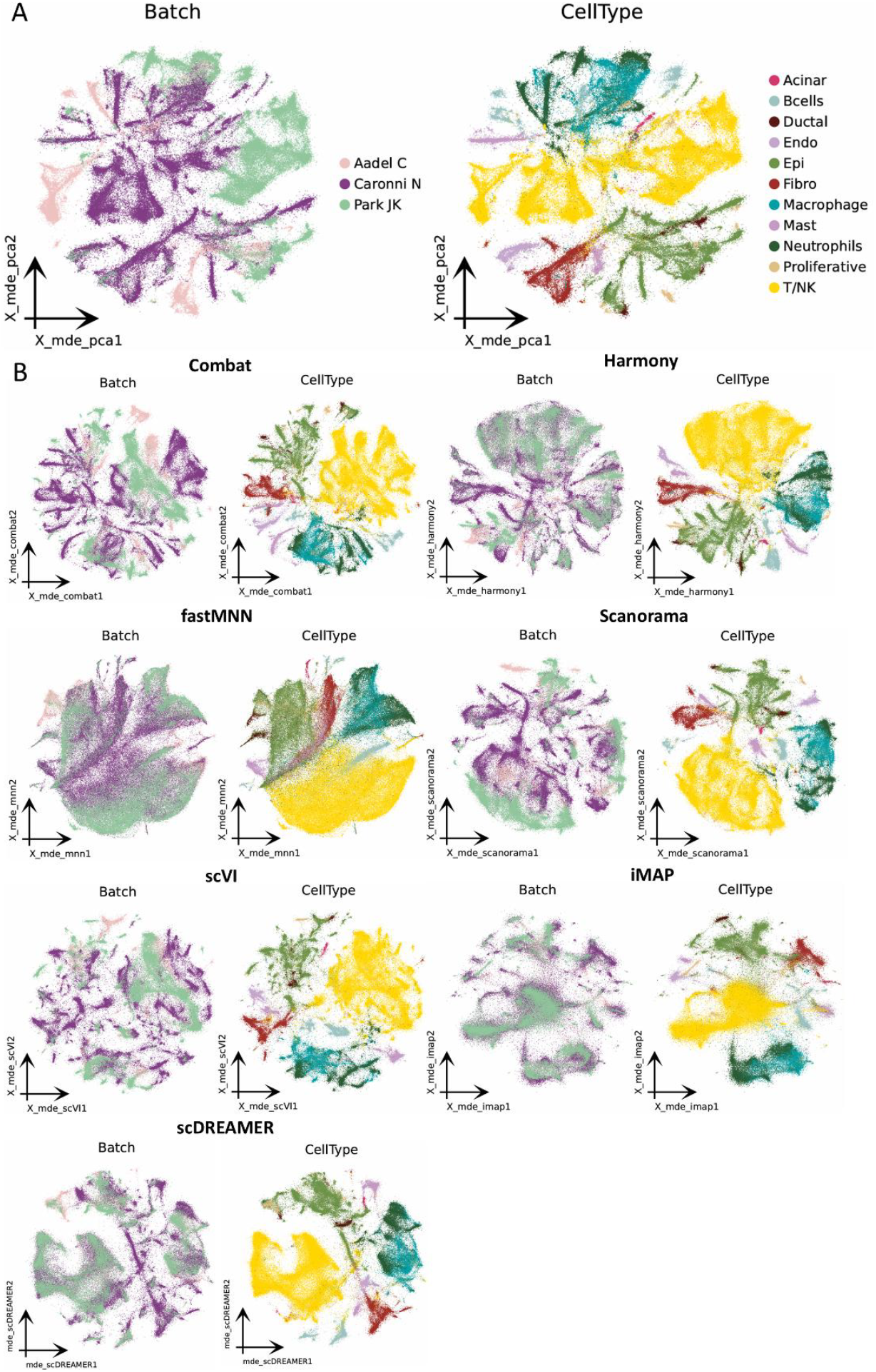
Integration of the PDAC dataset with mild batch effects. **(A)** MDE visualization of unintegrated data, colored by batch (left) and cell type (right). **(B)** MDE visualizations after integration by competing methods (ComBat, Harmony, fastMNN, Scanorama, scVI, iMAP), colored by batch.

**Fig. S2.**
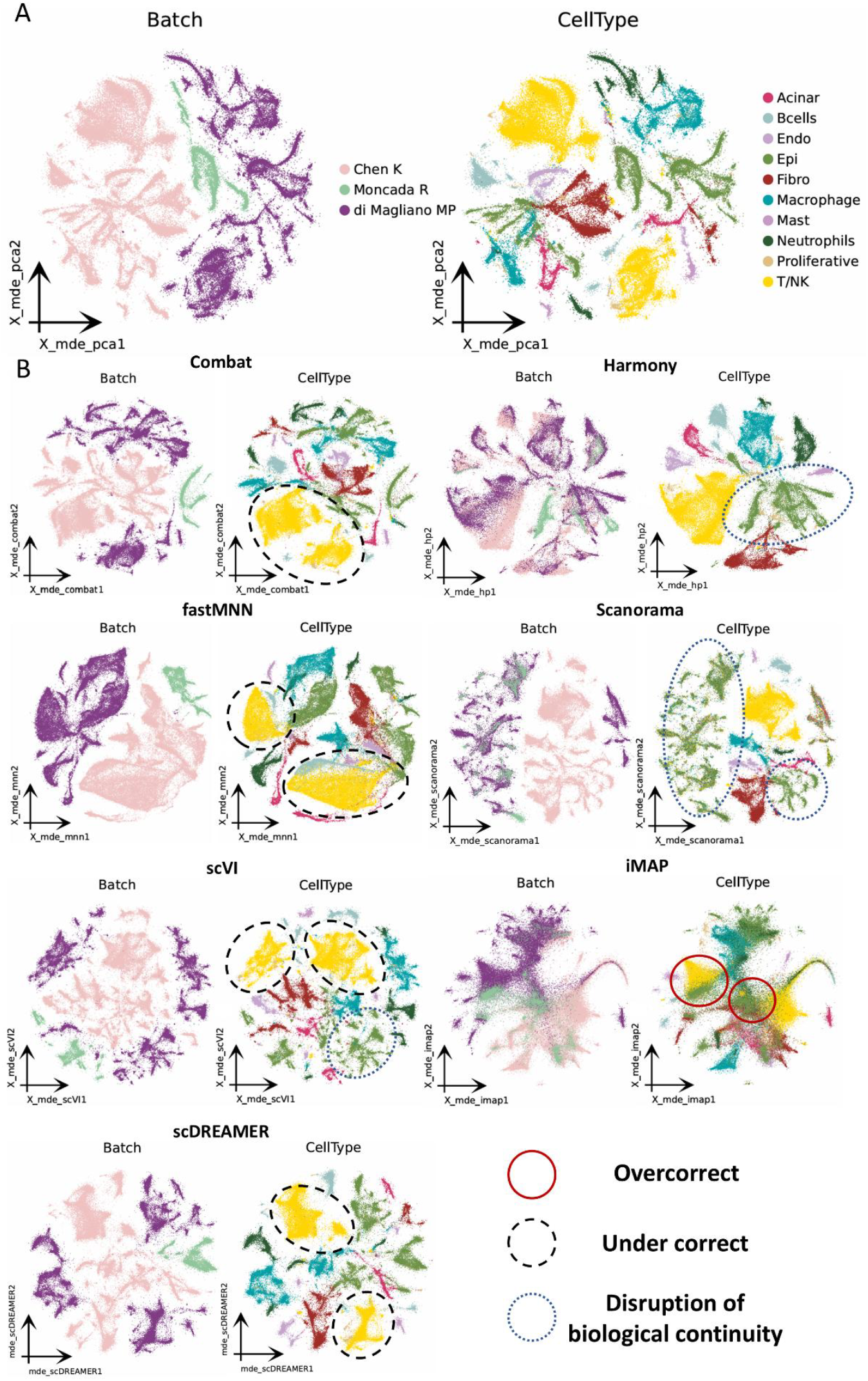
Integration of the PDAC dataset with strong batch effects. **(A)** MDE visualization of unintegrated data with strong batch effects. **(B)** MDE visualizations after integration by competing methods, colored by cell type, highlighting limitations such as insufficient mixing or over-correction.

**Fig. S3.**
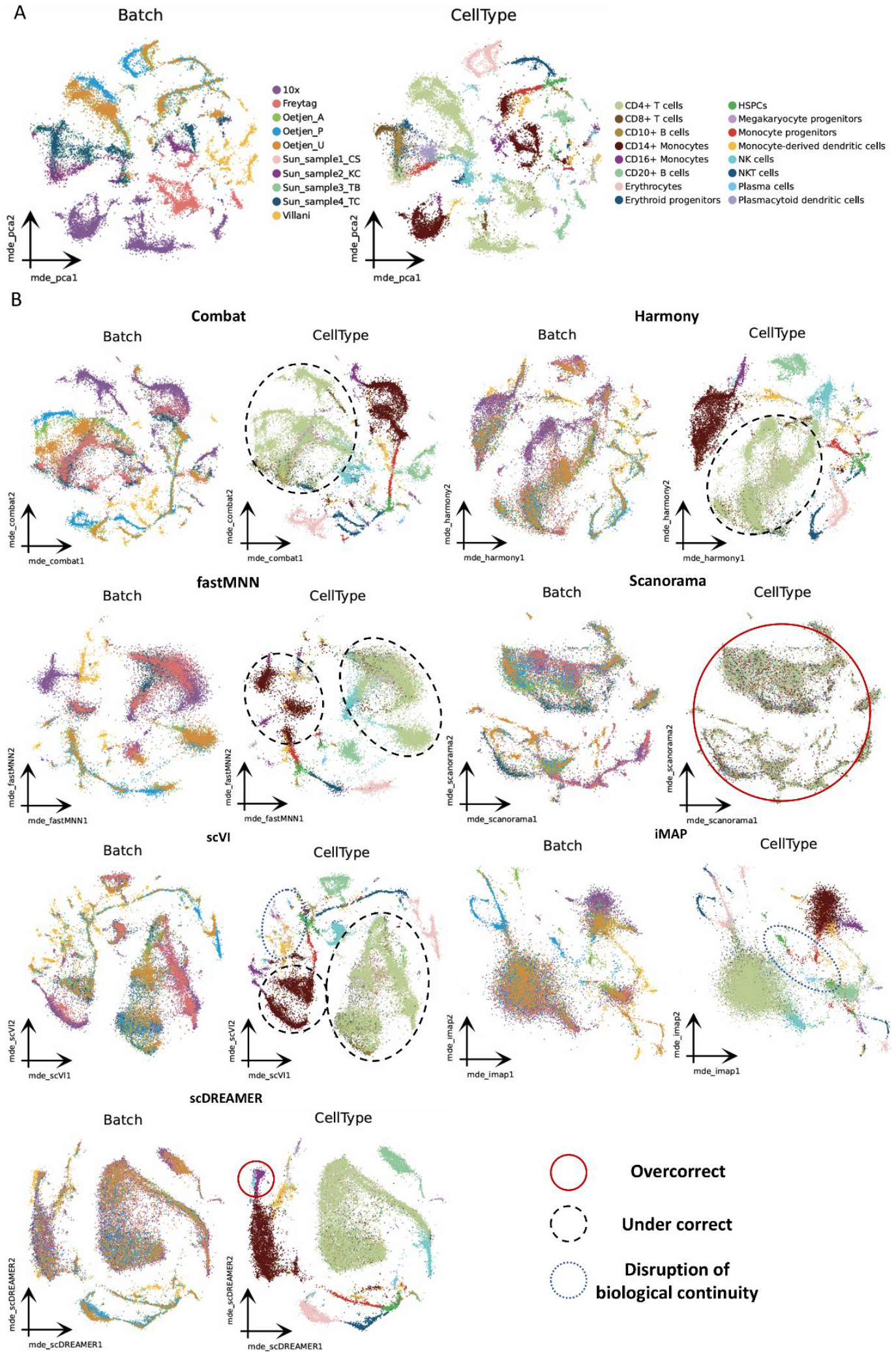
Integration of human immune cells across heterogeneous donors, tissues, and protocols. **(A)** MDE visualization of unintegrated human immune data. **(B)** MDE visualizations after integration by competing methods, colored by batch and cell type, revealing specific shortcomings in batch mixing and biological resolution.

**Fig. S4.**
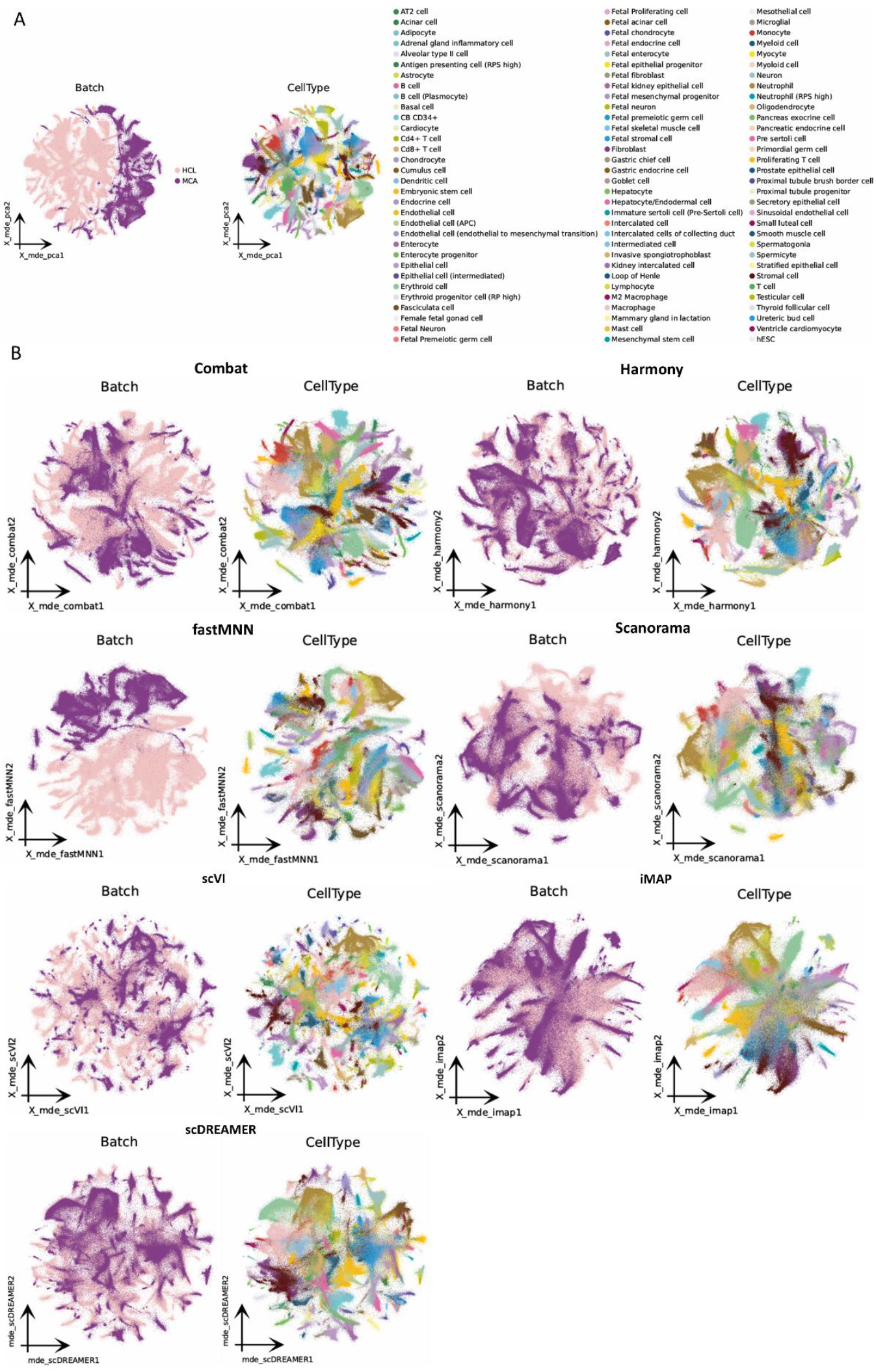
Integration of the cross-species human and mouse cell atlas data. **(A)** MDE visualization of unintegrated human and mouse cells, colored by species (left) and cell type (right).**(B)** Integrated embeddings from benchmark methods, colored by species and cell type. Results demonstrate common limitations in species mixing and structural preservation by existing approaches.

**Table S1.**
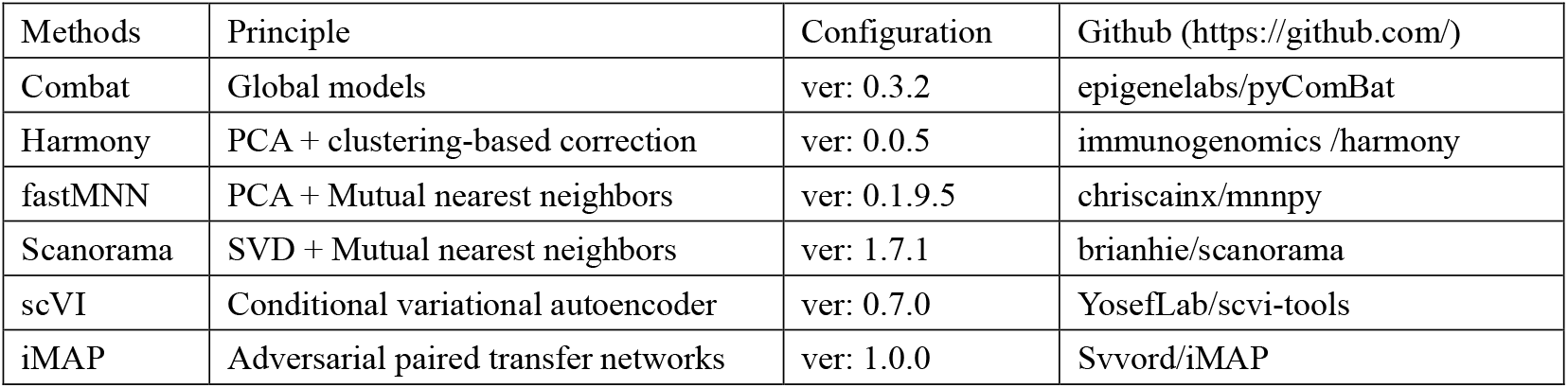
Configuration of the competing methods.

## Declarations

### Ethics approval and consent to participate

Not applicable

### Consent for publication

Not applicable

### Availability of data and materials

All datasets used in this study are publicly available. The human pancreatic ductal adenocarcinoma data used in this study are available in the GEO database under accession codes “GSE263733”, “GSE217845”, “GSE242230”, “GSE212966”, “GSE155698”, “GSE111672”. The human immune data used in this study are available in the GEO database under accession codes “GSE120221”, “GSE115189”, “GSE128066”, “GSE94820” and in the website of 10X Genomics (PBMC10k: https://support.10xgenomics.com/single-cell-gene-expression/datasets/3.0.0/pbmc_10k_v3). The Human and Mouse cell Atlas datasets used in the study are available at https://figshare.com/articles/dataset/MCA_DGE_Data/5435866 and https://figshare.com/articles/dataset/HCL_DGE_Data/7235471, respectively. The source code and usage tutorial for iDLC are freely available at https://github.com/zlsys3/iDLC.

### Authors’ contributions

S.L. and S.T. designed the study. C.J. developed models and algorithms, and wrote the manuscript. R.Z. and Y.F. further optimized the models. Y.J. and S.C. collected and annotated experimental data. Z.W. and R.W. reviewed and refined the manuscript. All authors have read and approved the final draft for submission.

## Acknowledgements

Not applicable.

